# The NOTCH3 Downstream Target HEYL Regulates Human Airway Epithelial Club Cell Differentiation

**DOI:** 10.1101/2021.03.10.434858

**Authors:** Manish Bodas, Bharathiraja Subramaniyan, Andrew R. Moore, Jordan P. Metcalf, Sarah R. Ocañas, Willard M. Freeman, Constantin Georgescu, Jonathan D. Wren, Matthew S. Walters

**Author notes:** **Correspondence:** Dr. Matthew S Walters, Assistant Professor, 800 N. Research Parkway, Building 800, 4th Floor, Rm 410, Oklahoma City, OK 73104, (on campus 13803).

## Abstract

Basal cells (BC) are the resident stem/progenitor cells of the adult pseudostratified airway epithelium, whose differentiation program is orchestrated by the NOTCH signaling pathway. NOTCH3 receptor mediated signaling regulates BC to club cell differentiation; however, the downstream responses that regulate this process are largely unknown. In the present study we used an *in vitro* air-liquid interface model of the human pseudostratified airway epithelium to identify the NOTCH3-dependent downstream genes/pathways that regulate human BC to club cell differentiation. Activation of NOTCH3 signaling in BC via lentivirus-mediated over-expression of the active NOTCH3 intracellular domain (NICD3) promoted club cell differentiation. Bulk RNA-seq analysis of control *vs* NICD3-transduced cells, identified 692 NICD3 responsive genes enriched for pathways linked to airway epithelial biology and differentiation including Wnt/β-catenin Signaling. Expression of the classical NOTCH target HEYL increased in response to NOTCH3 activation and positively correlated with expression of the club cell marker SCGB1A1. Further, using single-cell RNA-seq, we report that HEYL^+^ cells primarily clustered with SCGB1A1^+^ and NOTCH3^+^ cells. Moreover, HEYL protein co-localized with SCGB1A1 in ALI cultures *in vitro* and in the human and mouse airway epithelium *in vivo.* siRNA-mediated knockdown of HEYL in BC led to changes in epithelial structure including altered morphology and significant reductions in transepithelial electrical resistance and expression of tight junction related genes. Finally, HEYL knockdown significantly reduced the number of SCGB1A1^+^ club cells, along with a corresponding increase in KRT8^+^ BC-intermediate cells. Overall, our data identifies NOTCH3-HEYL signaling as a key regulator of BC to club cell differentiation.

## Introduction

The pseudostratified epithelium is a multi-cellular tissue that lines the conducting airways and functions as a barrier to protect the lung from environmental insults [1–4]. Basal cells (BC) are the resident stem/progenitor cells of the pseudostratified airway epithelium in humans and mice that initiate repair of the epithelium during homeostasis and following injury [1–9]. In response to specific stimuli, BC first differentiate into secretory club cells via BC-derived intermediate cells (also termed supra-BC or secretory primed-BC), which in turn undergo differentiation into either goblet secretory or ciliated cell lineages [5–9]. A reduction in the number of club cells is associated with human chronic lung disease, including asthma, chronic obstructive pulmonary disease (COPD) and idiopathic pulmonary fibrosis (IPF) [10–13]. Therefore, identifying the mechanisms that regulate BC to club cell differentiation are central to understanding the pathophysiology of chronic lung disease and the lung’s response to injury.

The NOTCH signaling pathway regulates BC stem/progenitor function and cell fate decisions in the human and murine airway epithelium [5, 14–44]. Canonical NOTCH signaling is initiated by the binding of a ligand (DLL1, DLL3, DLL4, JAG1 or JAG2) to one of the four receptors (NOTCH1-4) located on the surface of a neighboring cell. This leads to proteolytic cleavage of the receptor and release of the Notch intracellular domain (NICD) into the cytoplasm [45, 46]. The NICD then translocates to the nucleus and regulates expression of multiple downstream genes [45, 46]. Despite the knowledge that activation of NOTCH3 signaling is critical for BC to club cell differentiation [5, 19, 27], the downstream genes and pathways that regulate this process are unknown. The present study was designed to address this gap in our knowledge. Using the *in vitro* air-liquid interface (ALI) system to mimic the human pseudostratified airway epithelium, we have characterized the NOTCH3-dependent downstream genes/pathways in differentiating BC and identified the transcription factor HEYL as a NOTCH3 target that regulates club cell differentiation.

## Materials and Methods

### Primary human bronchial epithelial cell (HBEC) culture

Primary HBECs from normal, nonsmokers were purchased commercially (catalog number CC-2540, Lonza, Morristown, NJ, USA). The cells were maintained in BronchiaLife™ epithelial airway medium (BLEAM) (catalog number LL-0023, Lifeline^®^ Cell Technology, Oceanside, CA, USA) supplemented with Penicillin (100 I.U./ml) - Streptomycin (100 μg/ml) under standard cell culture conditions as previously described [14]. All experiments were performed with either Passage 2 or 3 cells. In total, n=9 cell donors were used in this study with the following lot numbers and demographics: Donor 1 (lot number: 0000501936, female, Hispanic, 42 years old), Donor 2 (lot number: 0000544414, male, Caucasian, 48 years old), Donor 3 (lot number: 0000619261, male, Caucasian, 53 years old), Donor 4 (lot number: 0000543643, female, Caucasian, 57 years old), Donor 5 (lot number: 0000613375, female, Black, 65 years old), Donor 6 (lot number: 0000529235, female, Black, 67 years old), Donor 7 (lot number: 0000608196, male, Caucasian, 67 years old), Donor 8 (lot number: 0000420927, female, Hispanic, 69 years old) and Donor 9 (lot number: 0000444771, male, Black, 69 years old).

### Air-liquid interface (ALI) culture

HBECs were differentiated using ALI culture to generate a pseudostratified airway epithelium as described previously [14]. Briefly, 1 × 10^5^ cells in 100 μ1 of BLEAM media were seeded in the apical chamber of a 0.4 μm pore-sized Transwell^®^ insert (catalog number 3470, Corning^®^, Corning, NY, USA) pre-coated with human type IV collagen (catalog number C7521, Sigma Aldrich, St. Louis, MO, USA) with 1 ml of BLEAM in the basolateral chamber (ALI day-2). The next day, fresh BLEAM media was replaced in the apical and basolateral chambers (100 μ1 and 1 ml, respectively). After two days of submerged culture, media from the apical chamber was removed to expose the cells to air (ALI day 0), and 1 ml of Human Bronchial/Tracheal Epithelial Cell (HBTEC) ALI differentiation medium (Cat# LM-0050, Lifeline^®^ Cell Technology) supplemented with Penicillin (100 I.U./ml) - Streptomycin (100 μg/ml) was added to the basolateral chamber. The media was replaced every other day and the cells allowed to differentiate for up to 9 days.

### RNA extraction, cDNA synthesis and quantitative PCR analysis

Total RNA was extracted via direct lysis of cells in the ALI well using the PureLink™ RNA mini kit (catalog number 12183018A, Thermo Fisher Scientific, Waltham, MA, USA) and included DNase treatment (catalog number 12185-010, Thermo Fisher Scientific) on the column to remove contaminating genomic DNA. Complementary DNA (cDNA) was generated from 100 ng of total RNA per sample using random hexamers (Applied Biosystems™ High Capacity cDNA Reverse Transcription Kit, catalog number 4374966, Thermo Fisher Scientific) and subsequent quantitative PCR (qPCR) analysis performed using iTaq™ Universal SYBR^®^ Green supermix (catalog number 1725124, Bio-Rad, Hercules, CA, USA) on the Bio-Rad CFX96 Touch™ RealTime PCR system. All samples were analyzed in duplicate with relative expression levels determined using the dCt method with Actin Beta (ACTB) as the endogenous control. The following PrimePCR^TM^ gene-specific primers were purchased from Bio-Rad: ACTB (qHsaCED0036269), SCGB1A1 (qHsaCID0018013), NOTCH3 (qHsaCID0006529), HEY1 (qHsaCED0046240), HEYL (qHsaCID0006092), NRARP (qHsaCED0048313), KRT5 (qHsaCED0047798), CLDN3 (qHsaCEP0032466), OCLN (qHsaCEP0041012), TJP1 (qHsaCIP0031627), TJP2 (qHsaCIP0026125), TJP3 (qHsaCIP0030881), PARD3 (qHsaCIP0039350) and PARD6B (qHsaCEP0051277), and the assays performed using the manufacturer’s recommend cycling parameters.

### Lentivirus-based overexpression of NICD3

Generation of Control (empty vector) or NICD3 expressing replication deficient lentiviruses was described previously [14]. Stocks of Lenti-Control and Lenti-NICD3 were titrated by qPCR (catalog number LV900, ABM^®^ good, Richmond, BC, Canada) and their infectivity confirmed by flow cytometry via quantification of GFP^+^ cells. Cells were infected at a multiplicity of infection of 50 (5 × 10^6^ viral genomes/1 × 10^5^ cells) at the time of seeding the cells on ALI to ensure that >90% of cells were infected (GFP^+^) with each virus. To aid virus infection, the BLEAM media was supplemented with 2 μg/ml of polybrene (catalog number TR-1003-G, Sigma Aldrich). The next day, fresh media was added to the wells and the standard ALI protocol was continued until the day of harvest.

### Bulk RNA sequencing (Bulk RNA-seq)

Control or NICD3-transduced cells (n=6 donors) were harvested as a function of time on ALI (day 0, 3, 5 and 7) and the genome-wide transcriptome changes in response to NICD3 expression assessed by bulk RNA-seq. Total RNA was extracted from each sample (described above) and bulk RNA-seq was performed on a NextSeq 500 Flowcell, High SR75 (Illumina, San Diego, CA, USA) following library preparation using the QuantSeq 3’ mRNA-Seq Library Prep Kit FWD for Illumina (Lexogen, Vienna, Austria). RNA-seq data processing followed the guidelines and practices of the ENCODE and modENCODE consortia regarding proper experimental replication, sequencing depth, data and metadata reporting, and data quality assessment (https://www.encodeproject.org/documents/cede0cbe-d324-4ce7-ace4-f0c3eddf5972/). Raw sequencing reads (in a FASTQ format) were trimmed of residual adaptor sequences using Scythe software. Low quality bases at the beginning or the end of sequencing reads were removed using sickle then the quality of remaining reads was confirmed with FastQC. Further processing of quality sequencing reads was performed with utilities provided by the Tuxedo Suite software. Reads were aligned to the Homo sapiens genome reference (GRCh38/hg38) using the TopHat component, then cuffquant and cuffdiff were utilized for gene-level read counting and differentially expression analysis. Genes that were significantly differentially expressed in response to NICD3 for at least one time point were determined using a threshold on the false discovery rate (FDR) adjusted p-value of 0.05 (FDR adjusted p<0.05). Ingenuity Pathway Analysis (IPA) (Qiagen, Redwood City, CA, USA) was used to identify molecular pathways altered in response to NICD3 using an unrestricted analysis.

### Single cell RNA sequencing (scRNA-seq)

HBECs from a single donor were cultured on ALI for 9 days and harvested for scRNA-seq analysis. To generate single cell suspensions, the cells were trypsinized for 3 mins and following neutralization passed through a Flowmi™ tip strainer (40 μm porosity, catalog number H13680-0040, SP Bel-Art, Wayne, NJ, USA). Cells were counted on a hemocytometer prior to diluting cells to 800 cells/μ1 in 0.1% BSA/PBS buffer for scRNA-seq library preparation with Chromium Single Cell 3’ Reagent Kits v3 (catalog number PN-1000075, 10X Genomics, Pleasanton, CA, USA). scRNA-seq libraries were generated according to 10X Genomics User Guide CG000183 Rev C. Briefly, a reaction mix containing 12,800 cells was loaded into a Chromium Chip B, targeting cell recovery of 8000 cells. After generating Gel Beads-in-emulsion (GEMS) on the Chromium controller, GEMS were transferred to PCR tubes (catalog number 951010022, Eppendorf, Hamburg, Germany) for GEM-Reverse Transcription (GEM-RT). After cleanup with Dynabeads MyOne Silane (catalog number PN-2000048, 10X Genomics), cDNA was amplified with 11 cycles. Amplified cDNA was cleaned with 0.6X SPRISelect reagent (catalog number B23318, Beckman Coulter, Pasadena, CA, USA). Cleaned cDNA was then quality checked on an Agilent Tapestation 4150 (catalog number G2992AA, Agilent, Santa Clara, CA, USA) using a High Sensitivity D5000 ScreenTape (catalog number 5067-5592, Agilent) and quantified using Qubit dsDNA High Sensitivity Assay Kit (catalog number Q32851, Thermo Fisher Scientific) read on a Qubit 4 Fluorometer (catalog number Q33238, Thermo Fisher Scientific). An aliquot of 25% of the cleaned cDNA was used for library construction (fragmentation, end repair, A-tailing, adaptor ligation), according to the manufacturer’s instructions. Libraries were indexed (catalog number PN-220103, 10X Genomics) using 10 cycles of PCR. Amplified libraries were cleaned with SPRISelect reagent using a double-sided size selection protocol. Cleaned libraries were quality checked on an Agilent Tapestation 4150 using a High Sensitivity D1000 ScreenTape (catalog number 5067-5584, Agilent) and quantified using Qubit dsDNA High Sensitivity Assay Kit. Libraries were diluted to 4nM prior to sequencing on a NovaSeq 6000 S1 flow cell with 28 cycles for read 1 and 90 cycles for read 2. Cellranger v3.1.0 (10X Genomics) cellranger mkfastq was used to demultiplex fastq files from raw base call (BCL) files. Fastq files were then aligned to Homo sapiens (human) genome assembly GRCh38-3.0.0 (hg38), filtered, and barcodes/UMIs counted using the cellranger count function. Summary metrics revealed 11,567 cells with an average of 8452 reads/cell and a median of 1382 genes/cell. scRNA-seq data was visualized using Loupe Cell Browser (10X Genomics).

### Data availability

The raw data from the bulk RNA-seq and scRNA-seq studies are publicly available at the Gene Expression Omnibus (GEO) site (http://www.ncbi.nlm.nih.gov/geo/), accession number GSE168128.

### siRNA mediated knockdown of NOTCH3 and HEYL

HBECs were transfected with either 1 pmol of *Silencer^®^Select* Negative Control No.1 siRNA (catalog number 4390844), NOTCH3 siRNA (catalog number 4392420; assay ID s453556), or HEYL siRNA (catalog number 4392420; assay ID s223702) using Lipofectamine RNAiMax Reagent (catalog number 13778075) and OptiMEM media (catalog number 31985070) (all from Thermo Fisher Scientific) at the time of seeding the cells in ALI culture. The following day, media in both apical and basolateral chambers was replaced with fresh BLEAM media, and the standard ALI protocol was continued until day of harvest (ALI day 7).

### Immunofluorescence staining and quantification of differentiation

Immunofluorescent staining of either differentiating cells in ALI wells or of the paraffin embedded human bronchus from healthy nonsmokers (Donor 1: Age 27, Female; Donor 2: Age 40, Female and Donor 3: Age 27, Female; catalog number HuFPT111, US Biomax, Inc., Rockville, MD, USA), or mouse trachea (C57BL/6, Normal, Female, 12 weeks) sections was performed as described [14]. For mouse trachea, the sections were subjected to an additional 1 hour of blocking at room temperature, with 20 μg/ml Affinipure Fab fragment goat anti-mouse IgG (H+L) (catalog number 115-007-003, Jackson ImmunoResearch Laboratories, Inc. West Grove, PA, USA). The following primary antibodies were used: HEYL (5 μg/ml, catalog number H00026508-M03, Abnova, Taipei, Taiwan), SCGB1A1 (5 μg/ml, catalog number RD181022220-01, BioVendor LLC, Asheville, NC, USA), KRT5 (2 μg/ml, catalog number PA1-37974, Thermo Fisher Scientific) and KRT8 (5 μg/ml, catalog number NBP2-16094, Novus Biologicals, Centennial, CO, USA). Isotype matched mouse (catalog number 401402, BioLegend) or rabbit IgG (catalog number 02-6102, Invitrogen) were used as a negative control. To visualize antibody binding the following secondary antibodies were used: goat anti-mouse Alexa Fluor 488 (2 μg/ml, catalog number A11029, Thermo Fisher Scientific) and goat anti-rabbit Alexa Fluor 546 (2 μg/ml catalog number A11035, Thermo Fisher Scientific). The cell nuclei were counterstained with DAPI (1 μg/ml, catalog number 62248, Thermo Fisher Scientific). Images were taken using an Olympus BX43 upright fluorescent microscope (Olympus Corporation). The number of cells positive for each marker of interest was counted using the ImageJ software (version 1.8.0_112, NIH), and normalized to the number of nuclei as described [14].

### Transepithelial electrical resistance (TEER)

TEER was measured using the ENDOHM-6G and EVOM2 apparatus (World Precision Instruments, Sarasota, FL, USA) according to the manufacturer’s guidelines. The resistance (ohms) of an empty Transwell^®^ insert (with no cells) was subtracted from each sample to calculate the true tissue resistance.

### Statistics

Statistical analysis was performed by two-tailed Mann Whitney U test using the GraphPad Prism version 8.0. A p value of ≤ 0.05 was considered significant.

## Results

To identify downstream genes and pathways that regulate NOTCH3-dependent differentiation of BC into club cells, bulk RNA-seq was performed on HBECs infected with control lentivirus or lentivirus expressing the constitutively active NICD3 as a function of time on ALI culture (day 0, 3, 5 and 7) (Fig. 1A-B). Quantitative PCR analysis of the club cell marker SCGB1A1 demonstrated increased club cell differentiation as a consequence of NICD3 over-expression in each HBEC donor (Fig. 1C). Comparison of NICD3-overexpressing *vs.* control cells identified 2319 genes with a significant (FDR adjusted p<0.05) expression change for at least one time point. For discovery purposes we increased the statistical stringency (FDR adjusted p<0.005) which reduced our list to 692 differentially expressed genes in response to NICD3 expression (Supplemental File 1). Analysis of the 692 gene set identified enrichment of molecular pathways including, “Hepatic Fibrosis / Hepatic Stellate Cell Activation”, “Inhibition of Matrix Metalloproteases”, “NOTCH Signaling”, “Integrin Signaling” and “Wnt/β-catenin Signaling” (Fig. 1D and Supplemental File 2). Expression of genes from each of these pathways correlated either positively or negatively with expression of SCGB1A1 in response to NICD3 expression (Fig. 1E). Further confirming the specificity of these downstream responses, siRNA knockdown of NOTCH3 (>70% knockdown efficiency in each experiment) led to an opposite effect to NICD3 over-expression, and decreased the expression of SCGB1A1 (0.38 fold) and the classical NOTCH downstream targets HEY1 (0.51 fold), HEYL (0.39 fold) and NRARP (0.40 fold) (Fig. 1F).

**Fig. 1.**
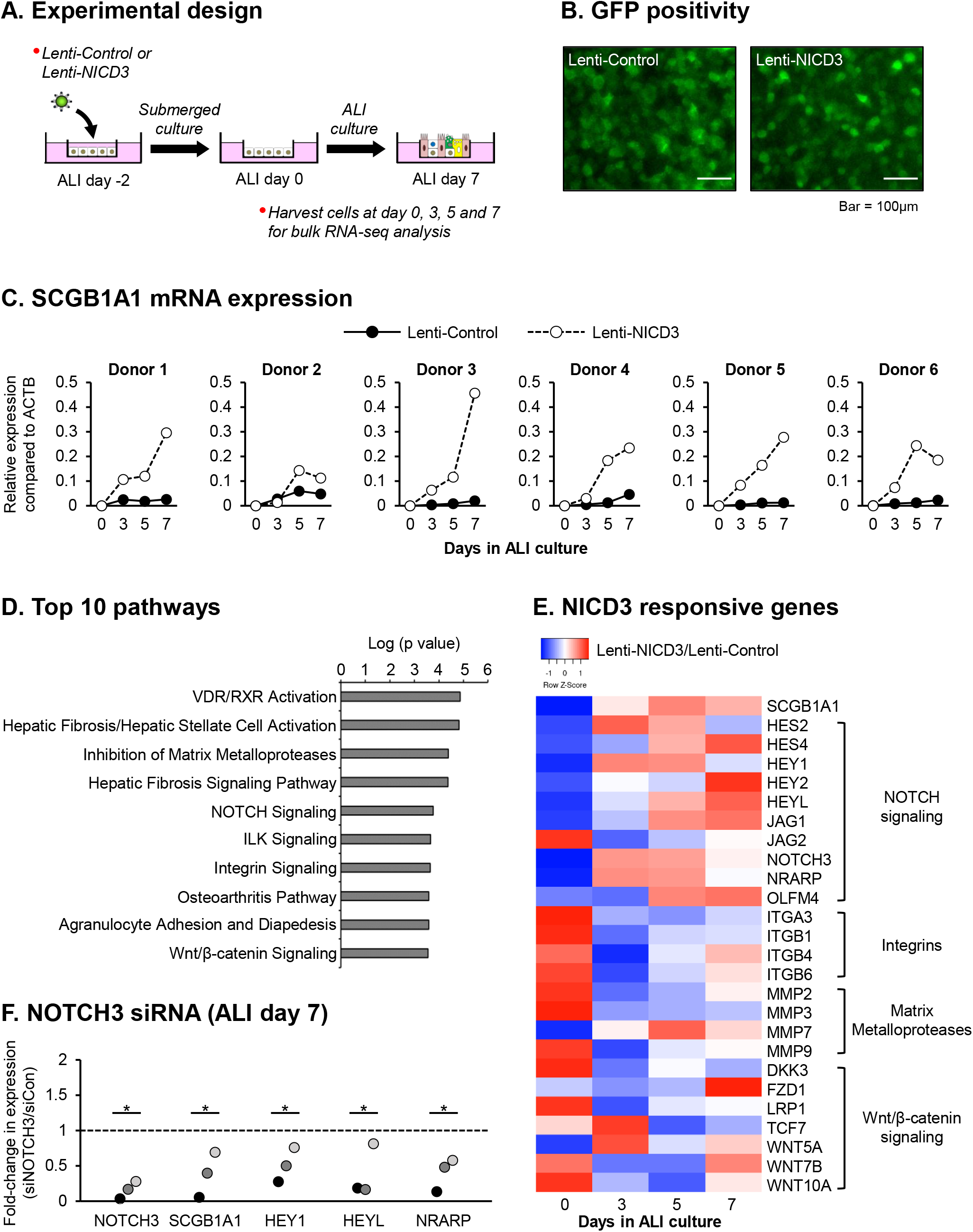
Activation of NOTCH3 signaling promotes club cell differentiation. **(A)** Experimental strategy. Primary HBECs (n=6 donors) were infected on air-liquid interface (ALI) with either control lentivirus (Lenti-Control) or lentivirus expressing the activated NOTCH3 NICD (Lenti-NICD3). The cells were then harvested as a function of time (ALI day 0, 3, 5 and 7) for subsequent analysis by qPCR and bulk RNA-seq. **(B)** GFP positivity of Lenti-Control and Lenti-NICD3 infected cells. Scale bar = 100μm. **(C)** qPCR of SCGB1A1 (club cell marker) expression in Lenti-Control and Lenti-NICD3 infected cells from each HBEC donor. **(D)** Pathways enriched in the 692 NICD3-responsive gene list identified by bulk RNA-seq on the basis of Ingenuity Pathway Analysis (IPA). Shown are the top ten IPA-enriched pathways based on P value (log-transformed). **(E)** Heatmap analysis of specific genes from the pathways highlighted in panel D. **(F)** HBECs (n=3 donors) were either transfected with control (siCon) or NOTCH3 (siNOTCH3) specific siRNA during seeding on ALI culture. At ALI day 7 the cells were harvested for qPCR analysis of NOTCH3, SCGB1A1, HEY1, HEYL and NRARP. For each donor, the data is presented as fold-change in expression compared to siCon cells. * p<0.05.

HEYL expression is upregulated in the murine tracheal airway epithelium following injury [32, 34]. However, the function of HEYL in the context of human lung differentiation is unknown. Quantitative PCR analysis showed that HEYL expression increased as a function of time in tandem with SCGB1A1 and NOTCH3 during HBEC differentiation on ALI (Fig. 2A). Furthermore, despite the low read depth limiting the detection of HEYL expression, scRNA-seq analysis of ALI day 9 cultures demonstrated that HEYL^+^ cells predominantly grouped with cells positive for SCGB1A1 and NOTCH3 in clusters 1 and 2 (Fig. 2B-F). Finally, co-immunofluorescent staining of *in vitro* ALI cultures (Fig. 2G) and *in vivo* human bronchial (Fig. 3A) and mouse tracheal epithelium (Fig. 3B) demonstrated that HEYL localizes with SCGB1A1^+^ club cells. Combined, these data suggest that induction of HEYL expression may regulate differentiation of BC into club cells. To test this hypothesis, HBECs were transfected with either control siRNA or HEYL specific siRNA and then cultured for 7 days on ALI. Compared to siRNA Control transfected cells, HEYL expression was significantly suppressed (0.06 fold) in siRNA HEYL transfected cells (Fig. 4A). Knockdown of HEYL led to changes in epithelial structure including altered morphology (Fig. 4B) and significant reductions in TEER (Fig. 4C) and expression of tight junction related genes (Fig. 4D). Furthermore, knockdown of HEYL significantly impaired club cell differentiation with a reduction in club cell numbers (0.53 fold) (Fig. 4E). While HEYL knockdown had no significant effect on the number of KRT5^+^ BC (Fig. 4F), we observed an increase in the numbers of KRT8^+^ BC-intermediate cells (2.42 fold) (Fig. 4G). Combined, these data suggest that HEYL is required for efficient differentiation of BC into club cells.

**Fig. 2.**
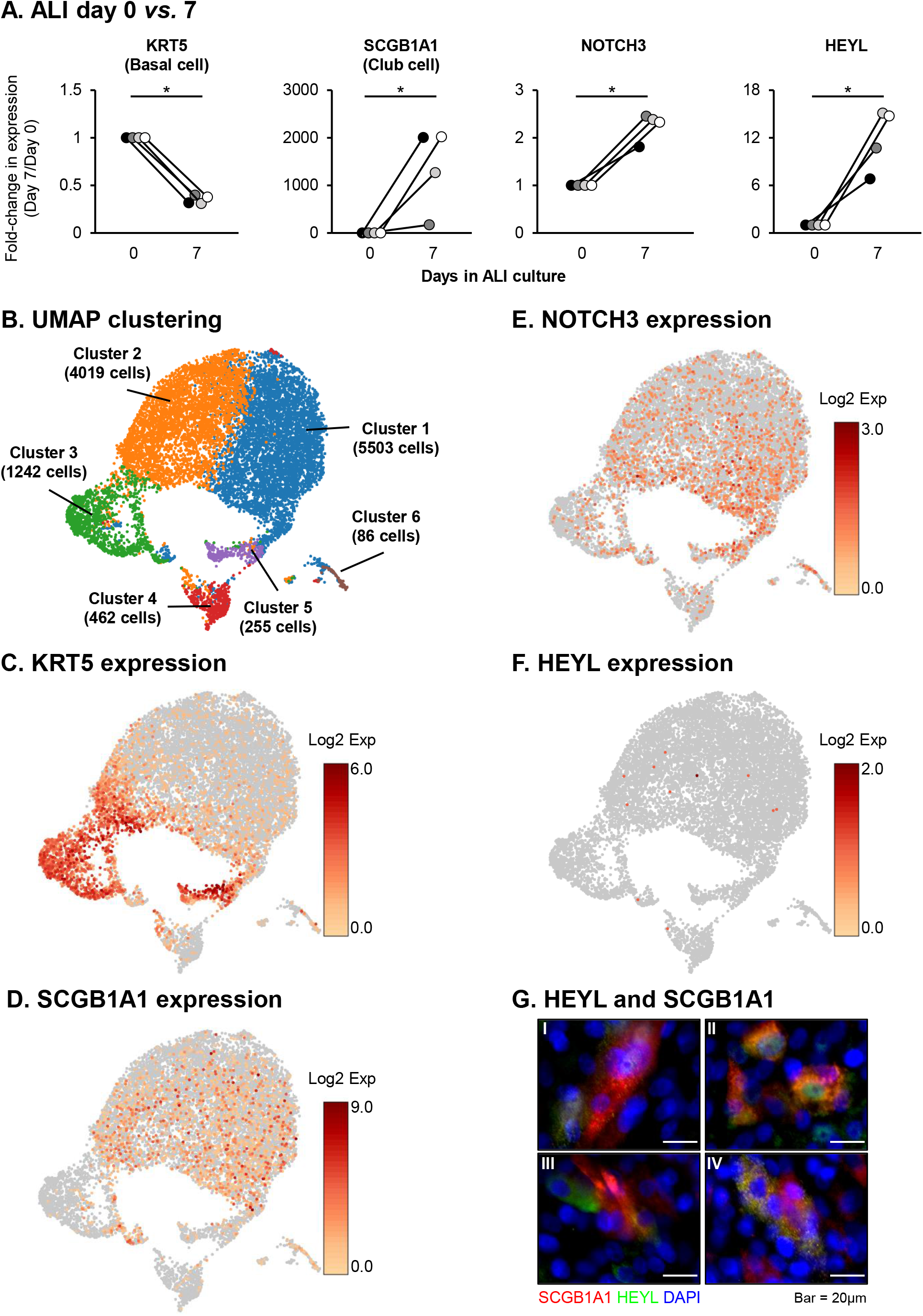
Expression of the NOTCH3 target HEYL correlates with club cell differentiation. **(A)** Primary HBECs (n=4 donors) were cultured on air-liquid interface (ALI) for 7 days. The cells were harvested at ALI day 0 and day 7 for qPCR analysis of KRT5 (basal cell marker), SCGB1A1 (club cell marker), NOTCH3 and HEYL. For each donor, the data is presented as fold-change in expression compared to ALI day 0. * p<0.05. **(B-F)** scRNA-seq analysis of HBECs cultured on ALI for 9 days. **(B)** UMAP clustering using K-means 6 clustering approach of 11,567 cells from a single HBEC donor. **(C)** Expression of KRT5 (basal cell marker). **(D)** Expression of SCGB1A1 (club cell marker). **(E)** Expression of NOTCH3. **(F)** Expression of HEYL. **(G)** Immunofluorescent staining of SCGB1A1 (red), HEYL (green) and nuclei (blue, DAPI) in ALI day 7 cells. Four representative images (I-IV) are shown. Scale bar = 20μm.

**Fig. 3.**
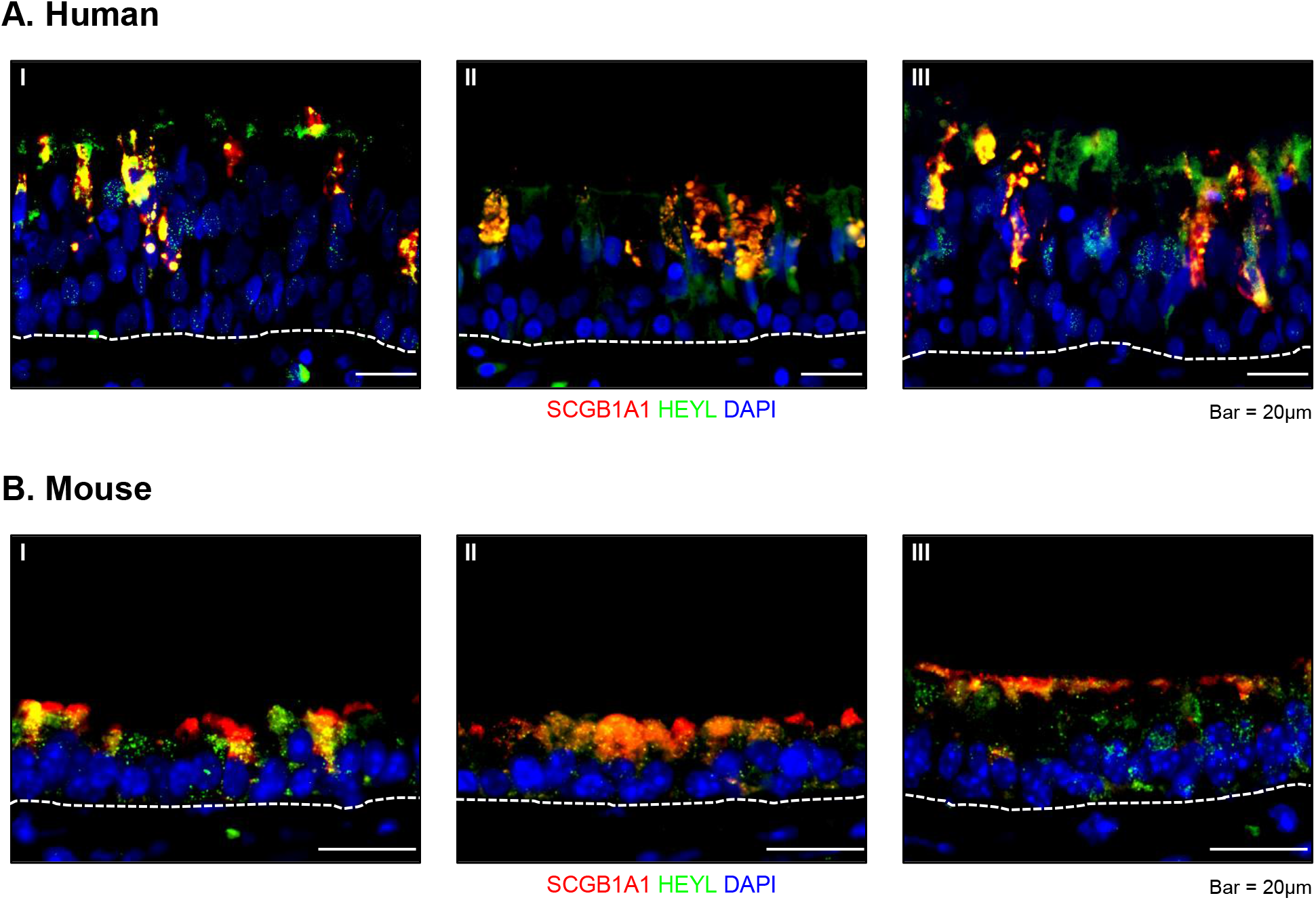
HEYL co-localizes with SCGB1A1 in the human and mouse airway epithelium *in vivo*. Immunofluorescent staining of SCGB1A1 (red), HEYL (green) and nuclei (blue, DAPI) in **(A)** human bronchus sections (n=3; donors) and **(B)** mouse trachea (n=3 mice). Scale bar = 20μm.

**Fig. 4.**
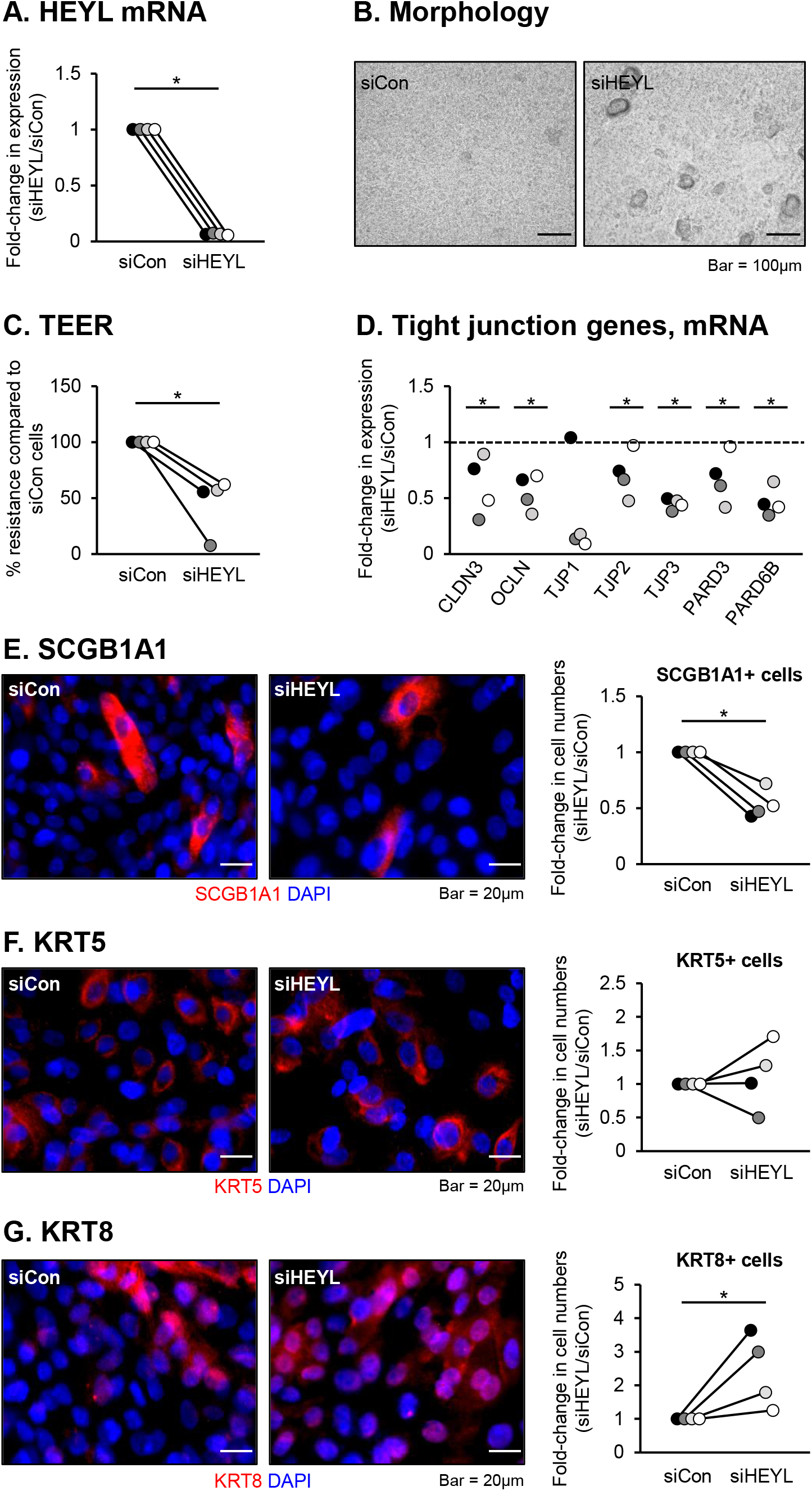
siRNA-mediated knockdown of HEYL suppresses club cell differentiation. Primary HBECs (n=4 donors) were transfected with either control (siCon) or HEYL (siHEYL) specific siRNA during seeding on ALI culture. At ALI day 7 the cells harvested for analysis. **(A)** qPCR analysis of HEYL. For each donor, the data are presented as fold-change in expression compared to siCon cells. * p<0.05. **(B)** Morphology of the cells following knockdown of HEYL expression. Scale bar = 100μm. **(C)** Transepithelial electrical resistance (TEER). For each donor, the resistance (ohms) is plotted as percentage (%) resistance compared to siCon cells. * p<0.05. **(D)** qPCR analysis of the tight junction related genes CLDN3, OCLN, TJP1, TJP2, TJP3, PARD3 and PARD6B. For each donor, the data are presented as fold-change in expression compared to siCon cells. * p<0.05. **(E-G)** Immunofluorescent staining and quantification of club cells (SCGB1A1, red), basal cells (KRT5, red) and basal-intermediate cells (KRT8, red). Nuclei are stained blue with DAPI. For each donor, the data are presented as fold-change in cell numbers compared to siCon cells. Scale bar = 20μm. * p<0.05.

## Discussion

NOTCH3 receptor signaling plays a key role in regulating differentiation of the pseudostratified airway epithelium during health and in chronic lung diseases, including asthma, COPD and IPF [5, 14, 19, 23, 27, 33]. To date, studies have shown that NOTCH3 signaling impacts airway epithelial differentiation in a context dependent manner with both suppression and activation of NOTCH3 signaling leading to pathological changes in airway epithelial structure [5, 14, 19, 23, 27, 33]. Therefore, greater understanding of the downstream genes and pathways that regulate NOTCH3-dependent differentiation are critical for understanding the mechanisms driving airway epithelial remodeling associated with chronic lung disease.

Based on the knowledge that NOTCH3 is an important regulator of club cell differentiation [5, 19, 27], we characterized the NOTCH3-dependent downstream genes/pathways in BC stem/progenitor cells as they differentiate into club cells on ALI culture. Bulk RNA-seq analysis identified 692 genes regulated in response to NOTCH3 activation that were enriched in molecular pathways associated with regulating airway epithelial structure and stem/progenitor cell function [24, 47–50]. These include “Hepatic Fibrosis/Hepatic Stellate Cell Activation”, “Inhibition of Matrix Metalloproteases” and “Integrin Signaling” [47, 48, 51]. Genes associated with these pathways play an important role in regulating the extracellular matrix (ECM), a structural scaffold that plays a critical role in regulating the growth, differentiation and function of airway epithelial cells [52, 53]. Furthermore, alterations in ECM structure is associated with the pathophysiology of multiple chronic lung diseases including asthma, COPD and IPF [52, 54]. Vera et al [55] recently reported a pathogenic role for NOTCH3 signaling in fibroblast activation and pulmonary fibrosis. Using a bleomycin-induced model of pulmonary fibrosis, they report that *Notch3*-deficient mice were protected from bleomycin-induced pulmonary fibrosis and had reduced numbers of αSMA-positive myofibroblasts compared to control fibrotic lungs. While mesenchymal cells (e.g., fibroblasts) are the major regulators of ECM deposition and organization, our data suggest that NOTCH3 signaling may have a similar role in regulating airway epithelial-derived ECM production. However, further studies are required to investigate the role of NOTCH3 in regulating these processes. Similar to NOTCH signaling, the Wnt/β-catenin signaling pathway also plays a crucial role in regulating cell fate decisions in the human lung [50, 56, 57]. Therefore, our data demonstrating NOTCH3 activation modulates expression of multiple genes in the Wnt/β-catenin signaling pathway suggests that NOTCH3 may cross-talk with this pathway to regulate airway epithelial differentiation.

At the single gene level, we also identified the transcription factor HEYL as a downstream target of NOTCH3 and a regulator of club cell differentiation. Expression of HEYL increases during differentiation on ALI culture in tandem with NOTCH3 and SCGB1A1 expression. Furthermore, HEYL is expressed in club cells in both the human (*in vitro* and *in vivo)* and mouse airway epithelium. Finally, knockdown of HEYL leads to a disruption in airway epithelial structure, as evident by decreased TEER values and reduced expression of tight junction genes, suggesting that HEYL directly regulates tight junction formation in the airway epithelium. Alternatively, the changes observed may be a consequence of decreased club cell differentiation and a corresponding increase in KRT8^+^ BC-intermediate cell numbers. Human studies have shown that HEYL regulates differentiation of fetal neural stem cells [58] as well as proliferation of breast, prostate and liver cancer cells [59–61]. Further, expression of HEYL is regulated in a NOTCH-independent manner by TGFβ signaling [59, 60], BMP signaling [62], and epigenetically via LSD1-mediated histone methylation [58] and DNA methylation of the promoter region [60, 63]. Expression of *Heyl* is upregulated in the murine tracheal airway epithelium following polidocanol and sulphur dioxide induced injury [32, 34], suggesting that *Heyl* may function in the regeneration response of the airway epithelium. However, this hypothesis has not been directly tested. Mori et al [27] reported that NOTCH3 signaling in the murine airway epithelium was critical for priming of BC differentiation into club cells and that *Notch3* knockout mice had increased numbers of KRT8^+^ undifferentiated progenitor cells in the airway epithelium compared to wild-type mice. These data support our findings whereby knockdown of HEYL leads to decreased club cell numbers and increased numbers of KRT8^+^ cells. Combined, these data suggest that NOTCH3/HEYL signaling is critical for regulating differentiation of BC into club cells and may be an early event in the regeneration response of the airway epithelium following injury. Therefore, future studies are required to explore the role of HEYL in regulating NOTCH3-dependent airway epithelial remodeling associated with human chronic lung disease.

## Acknowledgements

The authors thank Drs. Linda Thompson, Dean Dawson, Lorin Olson, and Xiao-Hong Sun at the Oklahoma Medical Research Foundation (OMRF) for discussions, guidance, and support. We also thank the Clinical Genomics Core and Imaging Core Facility at OMRF for assistance in sample processing related to our RNA-seq and histology studies.

## Declarations

### Ethics Approval

Not applicable.

### Consent to Participate

Not applicable.

### Consent for Publication

All the authors have provided consent for publication.

### Code availability

Not applicable.

### Author contributions

conception of study and experimental design: M.B. and M.S.W.; performed experiments: M.B., B.S., A.R.M., S.R.O., and M.S.W.; data analysis and/or interpretation: M.B., B.S., A.R.M., J.P.M., S.R.O., W.M.F., C.G., J.D.W. and M.S.W.; writing and preparation of original draft manuscript: M.B. and M.S.W. All authors have reviewed, critiqued, and approved the final manuscript.

